# Rapid inactivation *in vitro* of SARS-CoV-2 in saliva by black tea and green tea

**DOI:** 10.1101/2020.12.28.424533

**Authors:** Eriko Ohgitani, Masaharu Shin-Ya, Masaki Ichitani, Makoto Kobayashi, Takanobu Takihara, Masaya Kawamoto, Hitoshi Kinugasa, Osam Mazda

## Abstract

Saliva plays major roles in human-to-human transmission of the SARS-CoV-2. Recently we reported that black, green and oolong tea significantly inactivated SARS-CoV-2 within 1 min. Theaflavin-3,3’-di-gallate (TFDG), theasinensin A (TSA) and (-) epigallocatechin gallate (EGCG) were involved in the anti-viral activities. Here we examined how long period is required for the compounds to inactivate the virus. We also assessed whether tea inactivates SARS-CoV-2 diluted in human saliva. Treatment of SARS-CoV-2 with 500 μM TFDG or TSA for 10 sec reduced the virus titer to undetectable levels (less than 1/1,000). Black and green tea decreased virus titer to less than 1/100 within 10 sec even in saliva. These findings suggest a possibility that intake of, or gargling with, tea may inactivate SARS-CoV-2 in saliva in infected individuals, which may eventually attenuate spread of COVID-19 within a population, although clinical studies are required to test this hypothesis by determining the intensity and duration of the anti-viral effect of tea in saliva in humans.

## Introduction

The pandemic of novel coronavirus SARS-CoV-2 is expanding across the world with more than 80 million confirmed cases, and global death toll from COVID-19 has exceeded 1.7 million. Prophylactic procedures to attenuate the spread of virus infection is urgently needed.

SARS-CoV-2 is transmitted from a person to another principally through droplet infection, contact infection and aerosol infection ^1^. SARS-CoV-2 in saliva, sputum or nasal discharge is released from patients or infected individuals by talking, sneezing, coughing, and singing, and reaches nasopharyngeal, oral, or conjunctival mucosa of nearby persons through inhalation, ingestion, touching by fingers, etc ^2–6^. Especially, virus in saliva plays important roles, because saliva is commonly excreted during talking that is essential for our ordinary communication to each other as well as for economic and social activities in our societies. If the SARS-CoV-2 in saliva of infected individuals is effectively inactivated by a procedure, such a procedure could be useful for attenuating spread of the virus infection within a population.

We recently found that SARS-CoV-2 is significantly inactivated by a treatment with green tea, roasted green tea, oolong tea, and most remarkably, black tea ^7^. One min was sufficient for the inactivation of the virus by tea. As the active constituents, we found that (-) epigallocatechin gallate (EGCG) and theaflavin-3,3’-di-gallate (TFDG) that are bioactive compounds in green and black tea, respectively, and theasinensin A (TAS) that is a dimer of EGCG strongly inactivated SARS-CoV-2. These compounds significantly inhibited interaction between recombinant ACE2 and RBD of the Spike proteins, which may be, at least partially, involved in the mechanism underlying the anti-coronavirus effect of tea.

Therefore, tea could be used to inactivate SARS-CoV-2 in saliva in infected persons simply by ingesting or gargling with it. To assess the potential usefulness of tea for this purpose, we thought we should first know by *in vitro* analysis (i) how long period is required for the inactivation of SARS-CoV-2 by tea, and (ii) whether tea can inactivate the virus present in human saliva, before we start clinical studies to reveal (iii) whether tea ingestion/gargling inactivates the virus in human saliva *in vivo*, and (iv) how long period the viral inactivation effect in human saliva continues after tea ingestion/gargling.

In this study we tried to find answers to the above two questions (i) and (ii) in *in vitro* analyses.

## Results

We recently reported that treatment of SARS-CoV-2 with 500 μM of TSA or TFDG for 1 min significantly reduced infectivity of the virus ^7^. Then we treated the virus with 500 μM TSA and TFDG for various periods, and determined the virus titer by TCID_50_ assay (Fig. 1a). As results, 10 sec was enough for TSA and TFDG to reduce virus infectivity to below the detectable limit (less than 1/1,000) (Fig. 1b).

**Fig. 1.**
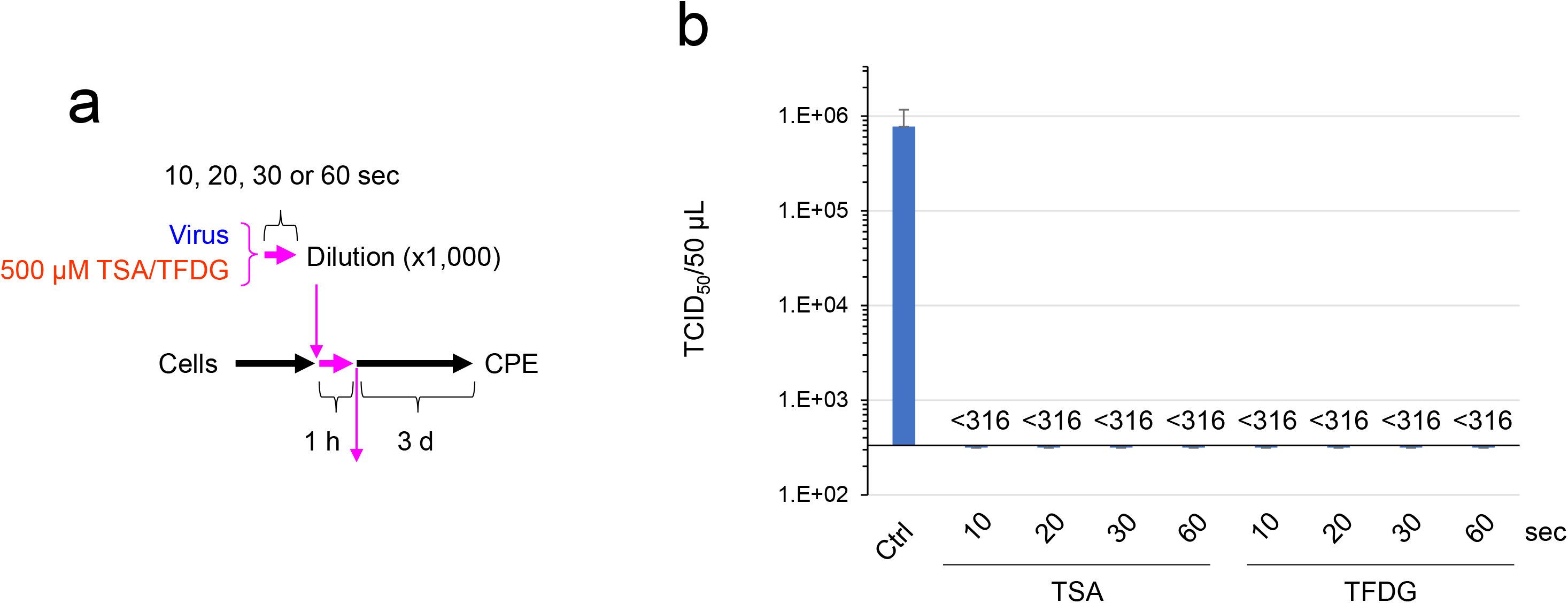
TSA and TFDG inactivated SARS-CoV-2 within 10 sec. SARS-CoV-2 was treated with 500 μM of TSA or TFDG for the indicated periods, immediately followed by a 1,000-fold dilution with MS. TCID_50_ assay was performed as described in the Materials and Methods. Scheme of experiment (a) and virus titer of each sample (means ± S.D., N=3) (b) are shown.

We tested whether black and green tea inactivated SARS-CoV-2 in human saliva. Virus suspension was diluted in saliva from five different donors (purchased from Lee Biosolutions)(Supplemenatry Table S1), and treated with 1 x black and green tea (vol:vol=1:1) for 10 sec (Fig. 2a). After dilution, virus was infected into cells. It was clearly shown that both black and green tea significantly declined the titer of the virus in saliva (Fig. 2b and c).

**Fig. 2.**
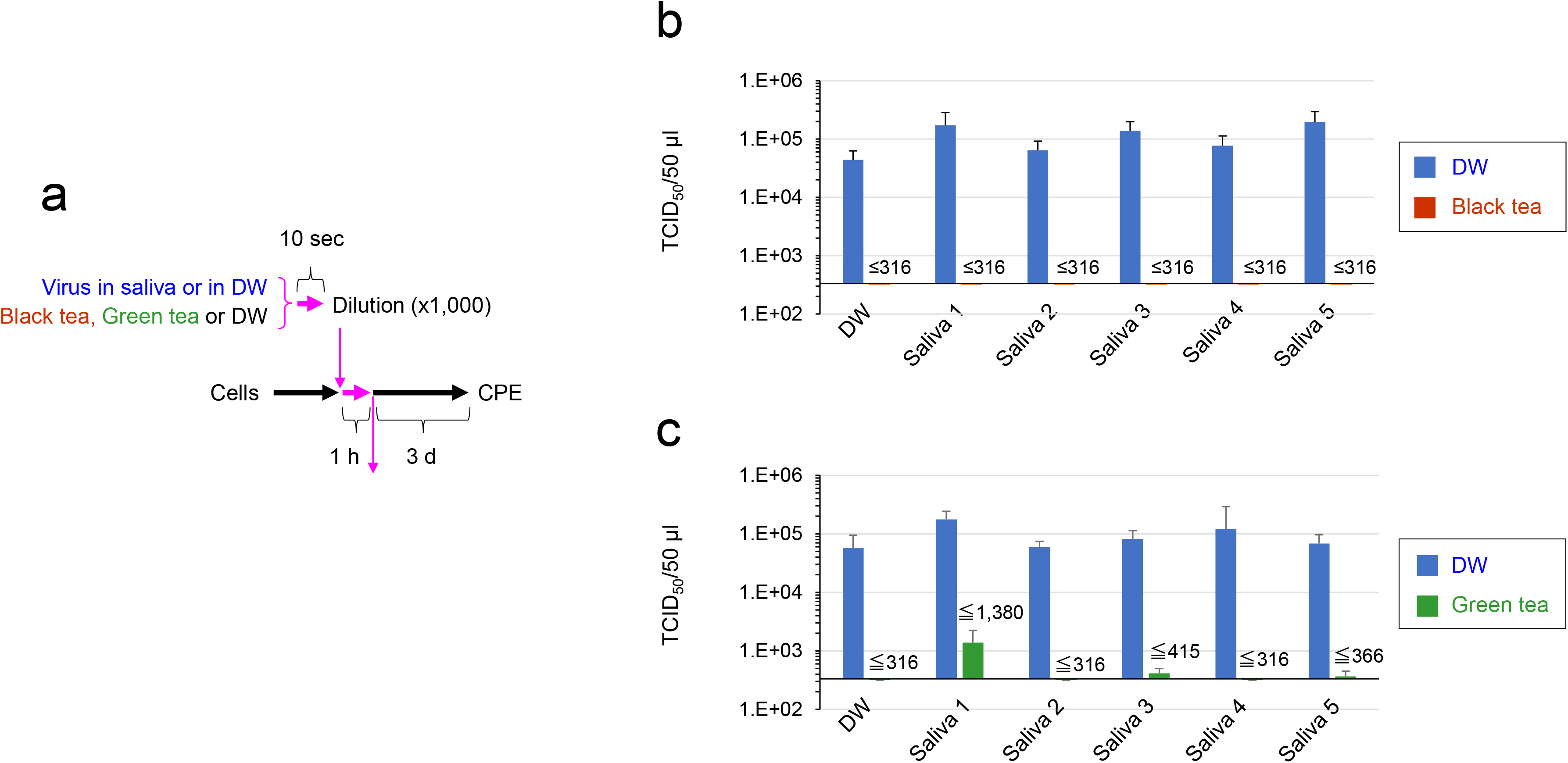
SARS-CoV-2 was inactivated by tea even in the presence of saliva. SARS-CoV-2 was diluted in saliva from five independent donors or in distilled water (DW) as a control (a). Black tea, green tea or DW was added to the virus/saliva for 10 sec, immediately followed by a 1,000-fold dilution with MS. TCID_50_ assay was performed as described in Fig. 1. Scheme of experiment (a) and virus titer of each sample (means ± S.D., N=3)(b) are shown.

To further confirm the virus inactivation caused by black tea treatment, virus in saliva was treated with black tea for 10 sec and infected to cells. After culturing, secondary virus generated from the cells was estimated (Fig. 3a). As shown in Fig. 3b, virus titers in culture supernatants were either not detected or significantly lower compared with the titer of secondary virus released from the cells infected with intact virus. The results strongly suggest that the SARS-CoV-2 treated with black tea replicated at a significantly low degree, if any, in cells even in the presence of saliva during the treatment.

**Fig. 3.**
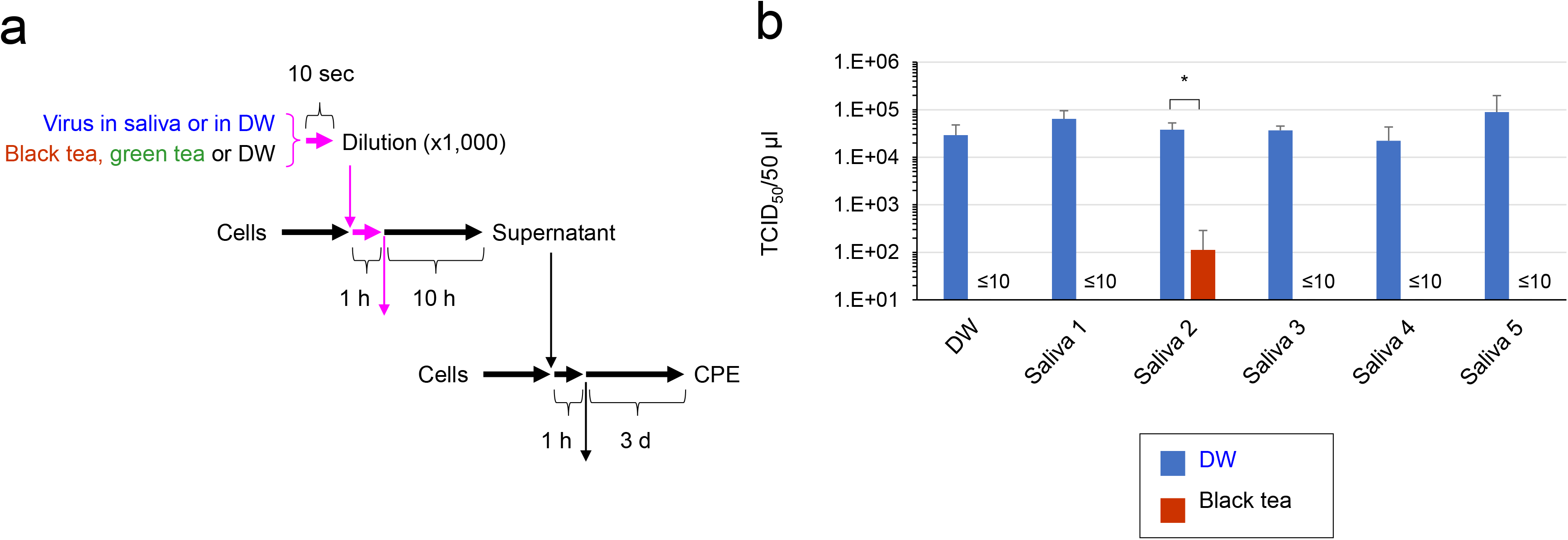
Treatment with tea significantly inhibited reproduction of SARS-CoV-2 in cells SARS-CoV-2 was diluted in saliva from five independent donors or in DW. Black tea or DW was added to the virus/saliva for 10 sec, immediately followed by a 1,000-fold dilution with MS. After infection, cells were cultured for 10 h, and culture supernatants were subjected to TCID_50_ assay as in Fig. 1. Scheme of experiment (a) and virus titer of each sample (means ± S.D., N=3)(b) are shown. *P<0.05 vs. DW.

## Discussion

Saliva is the major origin of droplets that are released from SARS-CoV-2-infected persons and cause droplet infection, contact infection and aerosol infection to nearby persons 6. Saliva contains various proteins including mucins, enzymes and immunoglobulins, nitrogenous products such as urea and ammonia, and electrolytes ^8^. The present study shows that tea significantly inactivates SARS-CoV-2 even in the presence of saliva (Figs. 2-4).

In the experiments shown in Figs. 2 and 3, the virus was treated with tea for 10 sec, and the mixture was immediately diluted 1,000-fold before an addition to the cells to allow viral attachment onto the cells. Tea diluted at 1,000-fold cannot effectively suppress virus infection as shown in our previous report ^7^. Therefore, only 10 sec was required for the inactivation of SARS-CoV-2 by tea.

Tea has a variety of biological effects including anti-viral activities, and tea catechins and their derivatives play important roles in these effects ^9–12^. But these compounds are poorly absorbed in the intestine ^13,14^, and their concentrations in plasma may not reach a sufficiently high level for inactivation of SARS-CoV-2. Therefore, tea may not be useful as a therapeutic medicine to treat respiratory and systemic manifestations of COVID-19 patients. Intake/gargling with tea by noninfected persons may be effective to some extent in preventing them from viral infection through the oral route, but may not be effective against viral entry via nasal and respiratory mucosa. But tea intake by SARS-CoV-2-infected persons could have greater significance as below.

It remains to be clarified to what extent SARS-CoV-2 in the saliva of SARS-CoV-2-infected persons can be inactivated *in vivo* by ingestion of, or gargling with, tea. Even if the virus in saliva is efficiently inactivated, the effect may be transient, because newly generated virus should reaccumulate in the saliva over time after the tea ingestion. Thus, it is important to reveal how long the anti-viral effect of tea will continue before virus reaccumulates in saliva to reach an enough high level to transmit infection to other persons through droplets. Turnover *in vivo* of the virus in saliva has not been reported so far, except for a study by Yoon et al. who examined a kinetic change of viral load in saliva in COVID-19 patients after mouthwash with chlorhexidine ^15^. They reported that the viral load dropped 1 hour after the disinfectant mouthwash in two patients. The viral load remained low for 2 hours, and jumped up at 3 hours, suggesting reaccumulation of newly generated virus between 2 and 3 hours after transient elimination of preexisting virus by the mouthwash.

If ingestion/gargling with tea results in remarkable inactivation *in vivo* of SARS-CoV-2 in saliva and the effect lasts for some hours, tea may be applied to a novel “public health intervention”-like approach to prevent virus transmission among people. Because there are a considerable number of asymptomatic carriers, a lot of people including “healthy” people should join the activity during a period of explosive surge of COVID-19, and ingest or gargle with a small amount of tea before they have close contact with other persons, talk, sing, play sports and music, etc. for a couple of weeks, not only to protect themselves but also to protect their neighbors. Such a “mutual defense” campaign based on altruistic behavior may potentially contribute to attenuation of spread of virus infection within a population. Besides, tea ingestion/gargling may be recommended to patients with other diseases who undergo dental treatment, upper GI endoscopic examination, and so on, to prevent nosocomial infection of SARS-CoV-2.

But if the anti-viral effect *in vivo* of tea is poor and/or continues only for a very short period, usage of tea may be limited. Therefore, careful clinical studies are needed to determine the efficacy and longevity of SARS-CoV-2 inactivation by tea in saliva of humans *in vivo*.

Finally, it should be noted that, although tea has been ordinary consumed by a large number of people for many years, intake of an excess amount of tea could cause adverse events including liver toxicity ^16,17^.

## Materials and Methods

### Cells, virus, and culture medium

VeroE6/TMPRSS2 cells ^18^ were obtained from Japanese Collection of Research Biosources Cell Bank, National Institute of Biomedical Innovation (Osaka, Japan) and cultured in Dulbecco’s modified Eagle’s minimum essential medium (DMEM) (Nissui Pharmaceutical Co. Ltd., Tokyo, Japan) supplemented with G418 disulfate (1 mg/mL), penicillin (100 units/mL), streptomycin (100 μg/mL), 5% fetal bovine serum at 37°C in a 5% CO_2_ / 95% in a humidified atmosphere. SARS-CoV-2 (Japan/AI/I-004/2020) were kindly provided from Japan National Institute of Infectious Diseases (Tokyo, Japan) and propagated using VeroE6/TMPRSS2 cells.

### Tea, chemicals and saliva

To prepare freeze-dried powders of tea extract, tea extracts were prepared by soaking 40 g of ground up and homogenized tea leaves in 2,000 mL water at 80°C for 30 min. After centrifugation at 4,000 rpm for 15 min, supernatants were collected and filtrated through Toyo No. 2 filter papers, followed by evaporation and freeze-drying. EGCG and TFDG were purchased from FUJIFILM Wako Pure Chemical Corporation (Osaka, Japan). TSA was synthesized from EGCG and purified as described. ^7,19^ Human saliva was purchased from Lee Biosolutions (Maryland Heights, MO, USA) (Supplementary Table 1).

### TCID_50_ assay for virus treated with TFDG and TSA

Solutions of TSA and TFDG were diluted in MS (DMEM supplemented with 0.5% FBS) to a concentration of 2 mM. Twenty μL of x2 serum-free DMEM was added to 20 μL of each solution, and 40 μL of SARS-CoV-2 suspension (2 × 10^6^ TCID_50_/50 μL) was added, followed by incubation at room temperature for 10, 20, 30 or 60 sec. Immediately, the mixture was diluted 1,000-fold in serum-free DMEM, followed by a serial dilution at 10-fold with MS in triplicate in 96-well-plates. Chilled on ice, 100 μL of each sample was added to the VeroE6/TMPRSS2 cells that had been seeded in 96-well-plates at 5 × 10^4^/100 μL/well a day before. After incubation for 1 hour to allow attachment of the virus onto the cells, supernatant was replaced by fresh culture medium, and cells were cultured for 4 days. Cells were then washed, fixed and stained with crystal violet solution to estimate CPE as described ^20^. TCID_50_ values were calculated by Reed-Muench method.

### TCID_50_ assay for virus in saliva treated with tea

Freeze-dried powders of green and black tea were dissolved in sterilized distilled water at 78°C for 10 min to prepare x1 concentration of original tea. After chilling at room temperature, each tea was passed through a 0.45 μm filter. Human saliva was sterilized by UV irradiation for 30 min. Virus suspension (3.0 × 10^5^ TCID_50_/5 μL) was mixed with 45 μL of saliva or water, followed by an addition of tea at 1:1 (vol:vol) for 10 sec. Immediately, the mixture was diluted 1,000-fold in serum-free DMEM, followed by a serial dilution at 10-fold with MS and TCID_50_ assay as above.

### TCID_50_ assay for secondary virus

Virus suspension in saliva was treated with black tea or distilled water for 10 sec, and immediately diluted 1,000-fold as above. One-hundred μL of the mixture was added to the cells that had been seeded in 24-well-plates at 2.5 × 10^5^/well a day before. After incubation for 1 hour to allow attachment of the virus onto the cells, supernatant was replaced by 500 μL of fresh culture medium, and cells were culture for 10 hours. An aliquot of culture supernatant was subjected to a serial dilution at 10-fold with MS and TCID_50_ assay as above.

### Statistical analysis

Statistical significance was analyzed by Student’s *t* test, and P<0.05 was considered significant.

---

**Supplementary Table S1.** Donors of saliva that was purchased from Lee Biosolutions (Maryland Heights, MO, USA) and used in the study are shown.

| Number | Birth year | Gender | Race |
| --- | --- | --- | --- |
| 1 | 1984 | Male | Caucasian |
| 2 | 1989 | Male | African American |
| 3 | 1986 | Male | Caucasian |
| 4 | 1967 | Female | Caucasian |
| 5 | 1968 | Male | Caucasian |

